# Designed peptides as potential fusion inhibitors against SARA-CoV-2 coronavirus infection

**DOI:** 10.1101/2020.06.09.142315

**Authors:** Ke Chen, Shihao Bai, Xin Zou, Jie Hao, Lin-tai Da, Ze-Guang Han

**Author notes:** To whom correspondence should be addressed, Ze-Guang Han. Correspondence may also be addressed to Lin-tai Da. or Jie Hao.

## Abstract

Inspired by fusion-inhibitory peptides from heptad repeat 1 (HR1) and heptad repeat 2 (HR2) domains from human immuno-deficiency virus type 1 (HIV-1) envelope glycoprotein gp41 and severe acute respiratory syndrome-coronavirus (SARS-CoV) based on viral fusogenic mechanism in the present work, we provided a similar approach to design the synthesized peptides against the entry into host cells of SARA-CoV-2 virus that causes 2019 novel coronavirus disease (COVID-19). These peptides derived from HR1 and HR2 of SARA-CoV-2 spike protein were further tested for their interaction and potential fusion possibility through circular dichroism spectrum. Here we used the peptide COVID-2019-HR1P1 (40 amino acids) as the target, which was derived from HR1 of SARA-CoV-2 spike protein, while the designed peptides including COVID-2019-HR2P1 (37 amino acids), COVID-2019-HR2P2 (32 amino acids) and others derived from HR2 of SARA-CoV-2 were tested for their binding to COVID-2019-HR1P1. Interestingly, results showed that both COVID-2019-HR2P1 and COVID-2019-HR2P2 can form the complex with COVID-2019-HR1P1, respectively. This implied that these designed peptides could play an important role in blocking SARA-CoV-2 entry into mammalian host cells via viral fusogenic mechanism, and thus could be used for preventing SARA-CoV-2 infection.

## INTRODOCTION

It has been just over 4 months since World Health Organization (WHO) announced the outbreak of new coronavirus SARS-CoV-2 infection as a Public Health Emergency of International Concern (PHEIC). By now, over 6.3 million cases are diagnosed as novel coronavirus pneumonia COVID-19 caused by SARS-CoV-2 infection, over 377 thousand people die worldwide.

Similar to severe acute respiratory syndrome-coronavirus (SARS-CoV) infection, the spike (S) protein of SARS-CoV-2 that leads to COVID-19 is considered to have strong binding affinity to the human cell receptor, angiotensin-converting enzyme 2 (ACE2)[1]. SARS-CoV-2 S protein (1273 amino acids) is consisted of two functional subunits: S1(aa 14-685) and S2(aa 686-1273) subunits. The former subunits is responsible for binding with ACE2 to form a S1-ACE2 complex, and then the whole S protein will be cleaved by host cell proteases into two separate fragments, S1 and S2. S2 subunit consists of a fusion peptide (aa 788-806), heptad repeat 1 (HR1) (aa 910-984) and heptad repeat 2 (HR2) (aa 1163-1213), which requires a 180°flip of prefusion helix fragments HR1 and HR2 to form a hairpin structure[2]. Because the fusion core regions are derived from HR1 and HR2 respectively, the two parts have strongly binding affinity with each other.

Previous study has shown the fragments COVID-2019-HR1P (aa 924-965), derived from HR1 of SARA-CoV-2 spike protein, and COVID-2019-HR2P (aa 1168-1203) from HR2 of SARA-CoV-2 are able to form a COVID-2019-HR1P-COVID-2019-HR2P complex[3]. In this study, we found more precise core binding regions to mediate HR1 and HR2 fusion as a viral fusogenic mechanism underlying the entry into host cells of SARA-CoV-2 virus.

## RESULTS AND DISCUSSION

### SARS-CoV and SARS-CoV-2 spike proteins and core fusion peptides

To evaluate whether SARS-CoV-2 virus employs the similar viral fusogenic mechanism for entering mammalian host cells, we compared SARS-CoV with SARS-CoV-2 spike protein sequences by BLAST tool. Interestingly, the known core fusion fragment (GVTQNVLYENQKQIANQFNKAISQIQESLTTTSTALGKLQ, 892-931) from HR1 of SARS-CoV has high similarity with COVID-2019-HR1P1 (GVTQNVLYENQKLIANQFNSAIGKIQDSLSSTASALGKLQ, 910-949) from HR1 of SARS-CoV-2 (Figure 1A and 1B), where 78% amino acids are identical among 40 amino acids between SARS-CoV and SARS-CoV-2. Moreover, the core fusion peptides (SARS-CoV-HR2P and COVID-2019-HR2P1) from HR2 of SARS-CoV and SARS-CoV-2 are 100% identical (Figure 1A). These facts suggest that, similar to SARS-CoV, SARA-CoV-2 virus could also enter host cells via a viral fusogenic mechanism.

**Figure 1.**
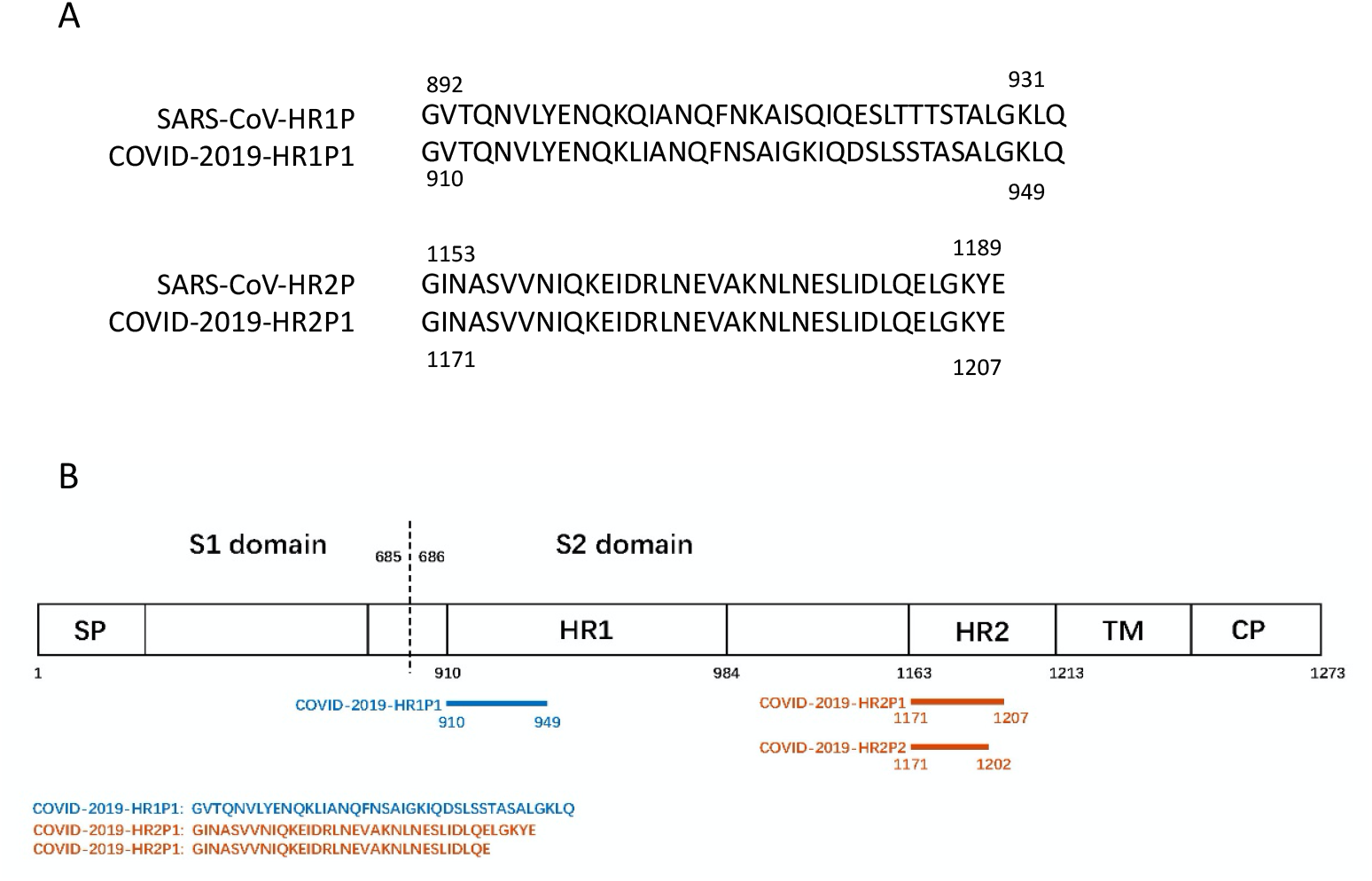
Schematic representation of SARS-CoV-2 spike protein and peptide sequences from HR1 and HR2 domains. (A) Spike protein sequences alignment with SARS-CoV and SARS-CoV-2 (labeled as COVID-2019). (B) Residue numbers of these examined peptide sequences correspond to their positions in SARS-CoV-2 spike protein. SP: signal peptide; HR: heptad repeat; TM: transmembrane domain; CP: cytoplasmic domain.

### Fusion analysis of SARS-CoV-2 HR1 and HR2 peptides based on circular-dichroism spectra

Based on above sequences analysis, 37 and 32 aa peptides (COVID-2019-HR2P1 and -HR2P2) from SARS-CoV-2 HR2 were synthesized and then tested the fusion possibility with HR1 of SARS-CoV-2 (COVID-2019-HR1P1) by Circular-dichroism spectra. Interestingly, the circular dichroism of their complex mixture have been modified (Figure 2). Here we noticed that the shorter peptide COVID-2019-HR2P2 has higher circular dichromatic absorption value than the longer peptide COVID-2019-HR2P1, although the latter also forms the complex with COVID-2019-HR1P1 based on the higher circular dichromatic absorption value. In order to acquire high fusion ability peptides, the mutants on COVID-2019-HR2P1 and COVID-2019-HR2P2 could be done in the future.

**Figure 2.**
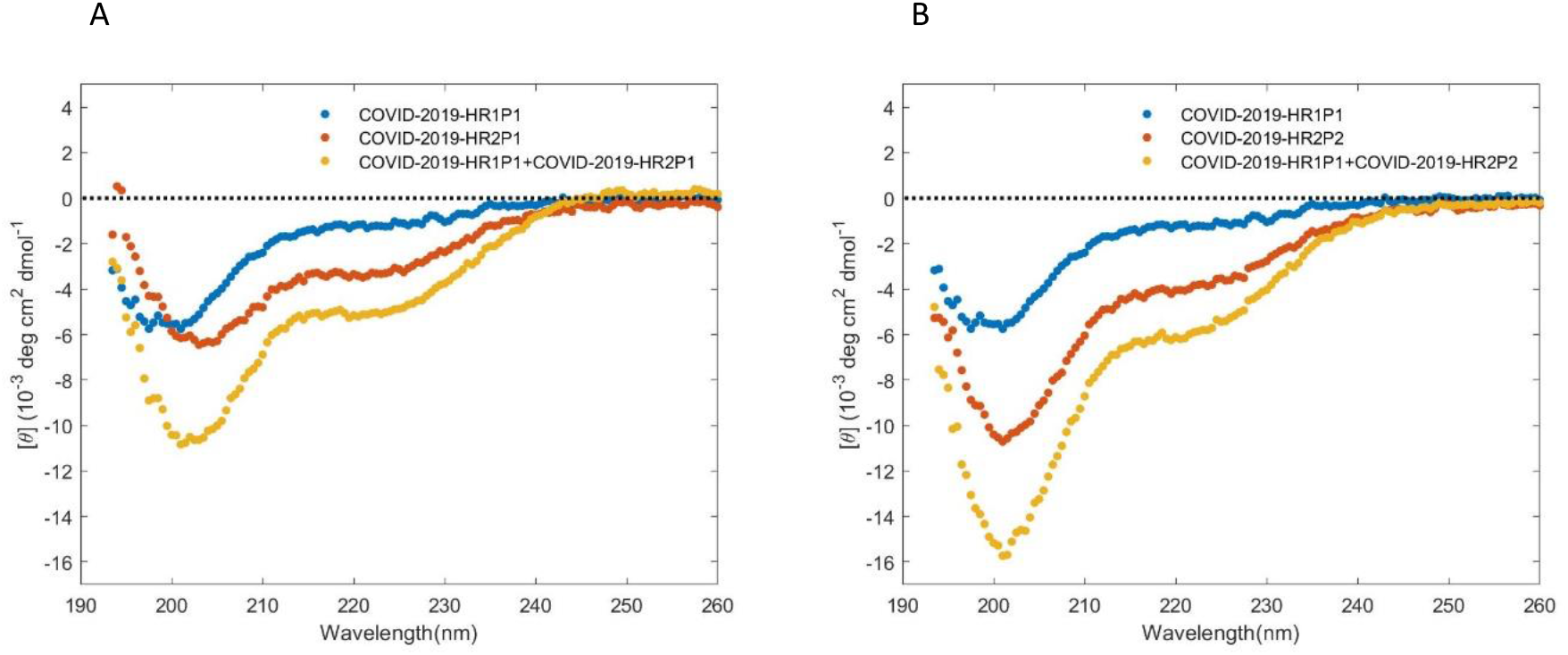
Circular-dichroism spectra reveals the fusion ability of these synthesized peptides derived from SARS-CoV-2 spike protein. (A) The fusion ability of the peptides COVID-2019-HR1P1 and COVID-2019-HR2P1. (B) The fusion ability of COVID-2019-HR1P1 and COVID-2019-HR2P2.

### Molecular modelling based on their secondary structure

To further reveal the fusion ability between the peptides, we predicted their interaction via calculating and analysing the molecular modelling among these peptides. Here one motif derived from HR2 of SARS-CoV-2, including G1171-E1207 (COVID-2019-HR2P1), was used as a potential fusion inhibitor via binding to the HR1 trimer (Figure 3). With an effort to enhance the inhibitory efficacy, several residue substitutions were introduced according to the structural features of the binding interface between HR1 (COVID-2019-HR1P1) and HR2 (COVID-2019-HR2P1). To pinpoint the key structural motif that contributes the most to the binding free energy, we also evaluated the inhibitory activities of one helical peptide from HR2 (COVID-2019-HR2P2). The result showed that both COVID-2019-HR1P1 and COVID-2019-HR2P1 as well as both COVID-2019-HR1P1 and COVID-2019-HR2P2 exhibited the strong interaction (Figure 3A and 3B). Here we got two different combined conformations, the peptides are aligned reversely or not (Figure 3A and 3B). Both corporation ways present similar pattern, where the amino acids distinct with SARS-CoV-HR1P always distribute away from the binding sites (Figure 3C), and however, the binding amino acids are totally nonpolar, including Val, Leu, Ile, Gly and Phe.

**Figure 3.**
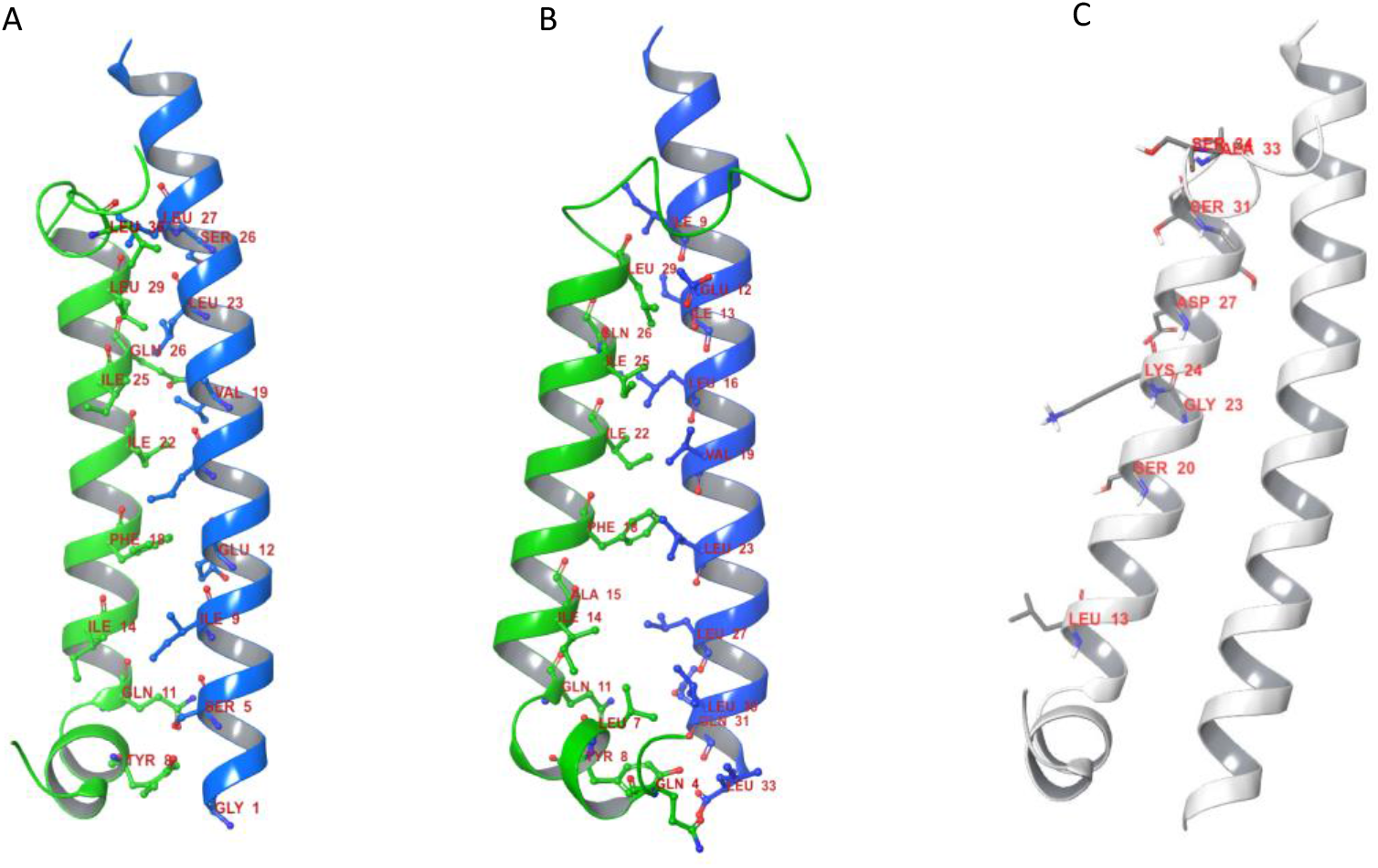
Interaction between these peptides as predicted by molecular modelling. (A) COVID-2019-HR1P1 (40 aa, green) and COVID-2019-HR2P1 conformation prediction (37 aa, blue) (binding reversely). (B) COVID-2019-HR1P1 and COVID-2019-HR2P2 conformation prediction (32 aa, blue) (syntropy binding). (C) Differential amino acids compared with SARS-CoV-HR1P are labelled.

## METHODS

### Circular-dichroism spectra detection

Circular-dichroism spectra for the peptides COVID-2019-HR1P1 (10 μmol/L), COVID-2019-HR2P1 (10 μmol/L), COVID-2019-HR2P2 (10 μmol/L) and their complexes were prepared in phosphate buffer (pH 7.2) at 4 °C.

### From sequence to 3D structure

COVID-2019-HR1P1 and COVID-2019-HR2P1 peptide sequences are imported to SWISS-MODEL (https://swissmodel.expasy.org)[4–8]. It builds the models by aligning with existing protein sequences before exporting the predicted 3D structure. We got the structure of COVID-2019-HR1P1 extracted from SARS-CoV-2 spike glycoprotein (Brookhaven Protein Databank, PDB ID:6VXX) and COVID-2019-HR2P1 from a trimer protein, 2019-nCoV HR2 Domain (PDB ID: 6LVN) (http://www.rcsb.org/).

### Interaction prediction on Maestro desktop

Maestro desktop from Schrodinger and OPLS-AA force field were used to do the pre-process work [9], including optimizing and minimizing. Water molecules and metal irons from proteins were removed. Then we chose peptide HR1P1 (40 amino acid) as rigid receptor, and modified the peptide HR2P1 7,000 times to get the top 15 stable structures which interpreted accurately as 15 clusters of structures. We got the median structure of the most stable cluster, along with one exceptional cluster, two proteins bind in the same direction.

